# iTraNet: A Web-Based Platform for integrated Trans-Omics Network Visualization and Analysis

**DOI:** 10.1101/2023.11.30.569499

**Authors:** Hikaru Sugimoto, Keigo Morita, Dongzi Li, Yunfan Bai, Matthias Mattanovich, Shinya Kuroda

## Abstract

A major goal in biology is to comprehensively understand molecular interactions within living systems. Visualization and analysis of biological networks play crucial roles in understanding these biochemical processes. Biological networks include diverse types, from gene regulatory networks and protein–protein interactions (PPIs) to metabolic networks. Metabolic networks include substrates, products, and enzymes, which are regulated by allosteric mechanisms and gene expression. Given this complexity, there is a pressing need to investigate trans-omics networks that include these various regulations to understand living systems. However, analyzing various omics layers is laborious due to the diversity of databases and the intricate nature of network analysis. We developed iTraNet, a user-friendly interactive web application that visualizes and analyzes trans-omics networks involving four major types of networks: gene regulatory networks (including transcription factor, microRNA, and mRNA); PPIs; metabolic networks (including enzyme, mRNA, and metabolite); and metabolite exchange networks (including transporter, mRNA, and metabolite). Using iTraNet, we found that in wild-type mice, hub molecules within the network tended to respond to glucose administration, whereas in *ob/ob* mice, this tendency disappeared. With its ability to facilitate network visualization and analysis, we anticipate that iTraNet will help researchers gain insights into biological systems. iTraNet is available at (https://transomics.streamlit.app/).

**GRAPHICAL ABSTRACT:** 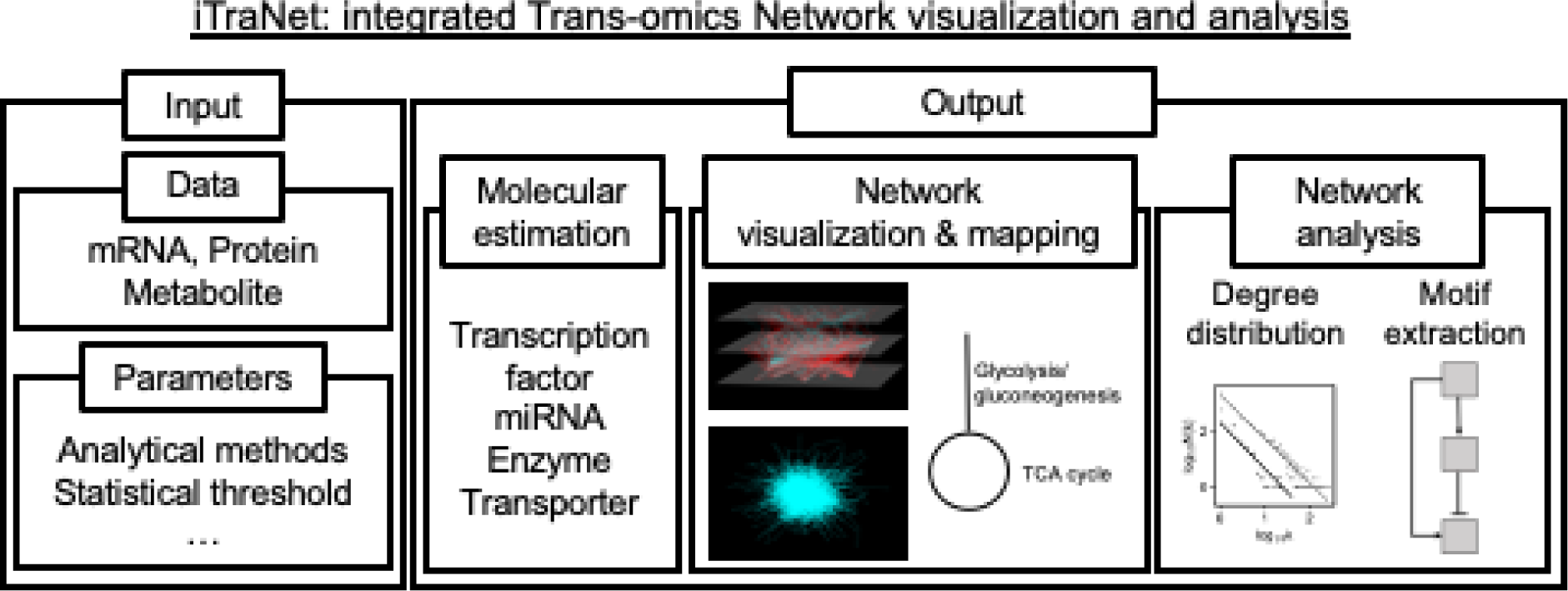

## INTRODUCTION

A major goal of biology is a comprehensive understanding of molecular interactions within biological systems (1). These interactions intricately regulate intracellular processes, leading to the emergence of biological network visualization and analysis as pivotal methodologies in biology (1–5). Through network visualization and analysis, researchers have a platform to explore relationships and interactions among molecules, revealing the underlying regulatory mechanisms that influence cellular behavior (2, 3).

One example of how analysis of the properties of entire biological networks has led to a better understanding of living systems is discovery of the robustness of “scale-free networks” to errors (6). Many complex systems, including metabolic networks, exhibit remarkable resilience to errors, with local failures rarely leading to a loss of global information transmission capacity within the networks (6). This error resilience is partly achieved by a class of network topology known as scale-free networks, characterized by heterogeneous wiring (6). Network analysis also guides hypothesis formation and the development of targeted intervention strategies by enabling the identification of pivotal nodes, hubs, and modules within complex networks (7–10).

Biological networks include many forms, ranging from gene regulatory networks and protein–protein interactions (PPIs) to metabolic networks. Metabolic networks encompass substrates and products in metabolic reactions, as well as the enzymes that direct these processes. These enzymes are subject to regulation via allosteric regulation and gene expression regulation. Gene expression is regulated by transcription factors (TFs) and microRNAs (miRNAs). In addition, intracellular processes interact with extracellular molecules via transporters. These rationales underscore the importance of applying multilayered trans-omics networks to understanding biological systems; we have applied this approach to understanding metabolism in living systems (3, 11–16).

Such a comprehensive understanding of living systems requires a comprehensive analysis of biological networks. However, analyzing various omics layers still remains a laborious and time-consuming task due to their derivation from heterogeneous databases and the necessity for complex network analysis. Ensuring the reproducibility of such studies is also challenging due to the continuously updating databases and occasional unavailability of publicly accessible codes for performing trans-omics analysis. Recognizing these problems, we developed a user-friendly and open access web application that performs trans-omics network visualization and analysis.

Here, we introduce iTraNet, a user-friendly web application designed to visualize and analyze trans-omics networks representing four key types of interactions in living systems: gene regulatory networks (including TF, miRNA, and mRNA); PPIs; metabolic networks (including enzyme, mRNA, and metabolite); and metabolite exchange networks (including transporter, mRNA, and metabolite). This web application not only visualizes the intricate network interplay among these molecules but also conducts some analyses of their network properties, providing researchers with a holistic understanding of intricate biological dynamics.

## MATERIALS AND METHODS

In this section, we summarize the approach that we followed to construct and analyze trans-omics networks and implement them as a web application. Further comprehensive details on the construction and analysis of trans-omics networks are available in our previous studies (3, 11–16).

### Workflow overview

The main aim of iTraNet is to provide researchers with an easily accessible platform for trans-omics network visualization and analysis. Figure 1 shows the overall workflow of iTraNet. To perform trans-omics network visualization and analysis in iTraNet, users are required to upload their own transcriptome, proteome and/or metabolome data (Fig. 1, left panel). Upon submission, iTraNet performs visualization and analysis across four distinct categories of biological networks: (A) gene regulatory networks (including transcription factor (TF), microRNA (miRNA), and mRNA); (B) protein (mRNA)-protein (mRNA) interactions; (C) metabolic networks (including enzyme, mRNA, and metabolite); and (D) metabolite exchange networks (including transporter, mRNA, and metabolite).

**Figure 1.**
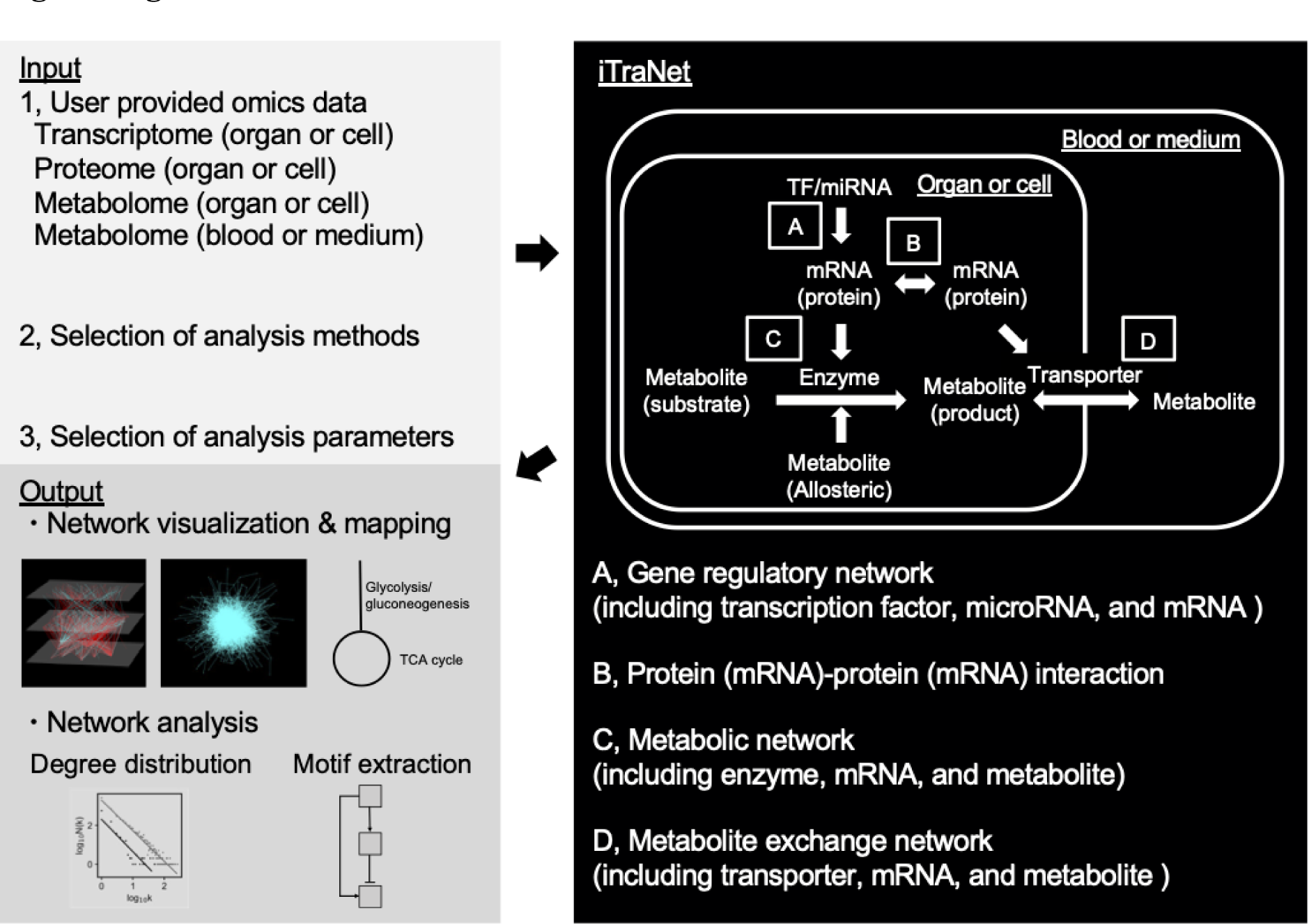
Schematic overview of iTraNet workflow. iTraNet requires transcriptome, proteome, and/or metabolome data, as input (left, top panel). iTraNet is designed to visualize and analyze four distinct categories of biological networks, as shown in the right panel: (A) gene regulatory networks (including transcription factors (TF), microRNA (miRNA), and mRNA); (B) protein (mRNA)–protein (mRNA) interactions; (C) metabolic networks (including enzyme, mRNA, and metabolite); and (D) metabolite exchange networks (including transporter, mRNA, and metabolite). Finally, iTraNet outputs a comprehensive visualization of the four types of biological networks, maps for each metabolic pathway, and results of the network analyses (left, bottom panel).

Estimating relationships within TF, miRNA, and mRNA networks (Fig. 1A) requires relevant transcriptome data from targeted organs or cells. Similarly, estimating relationships among mRNAs or proteins (Fig. 1B) necessitates transcriptome or proteome data from the targeted organs or cells. Meanwhile, assessing relationships involving enzymes, mRNAs, and metabolites (Fig. 1C) calls for both metabolome and transcriptome data from the targeted organs or cells. Estimating relationships among transporter, mRNA, and metabolite (Fig. 1D) requires metabolome and transcriptome data from the targeted organs or cells, as well as the associated blood or medium.

iTraNet estimates TFs and miRNAs associated with the uploaded mRNAs (Fig. 2A). The associated TFs and miRNAs are estimated using the ChIP-Atlas (17) and miRTarBase (18) databases, respectively. Users can set a threshold for statistical significance value calculated by MACS2 in estimating TFs, as previously described (17). Additionally, users can set a false discovery rate for TF enrichment analysis, performed using the one-tailored Fisher’s exact test followed by the Benjamini-Hochberg method. Mus musculus data in the miRTarBase was used for estimating miRNA.

**Figure 2.**
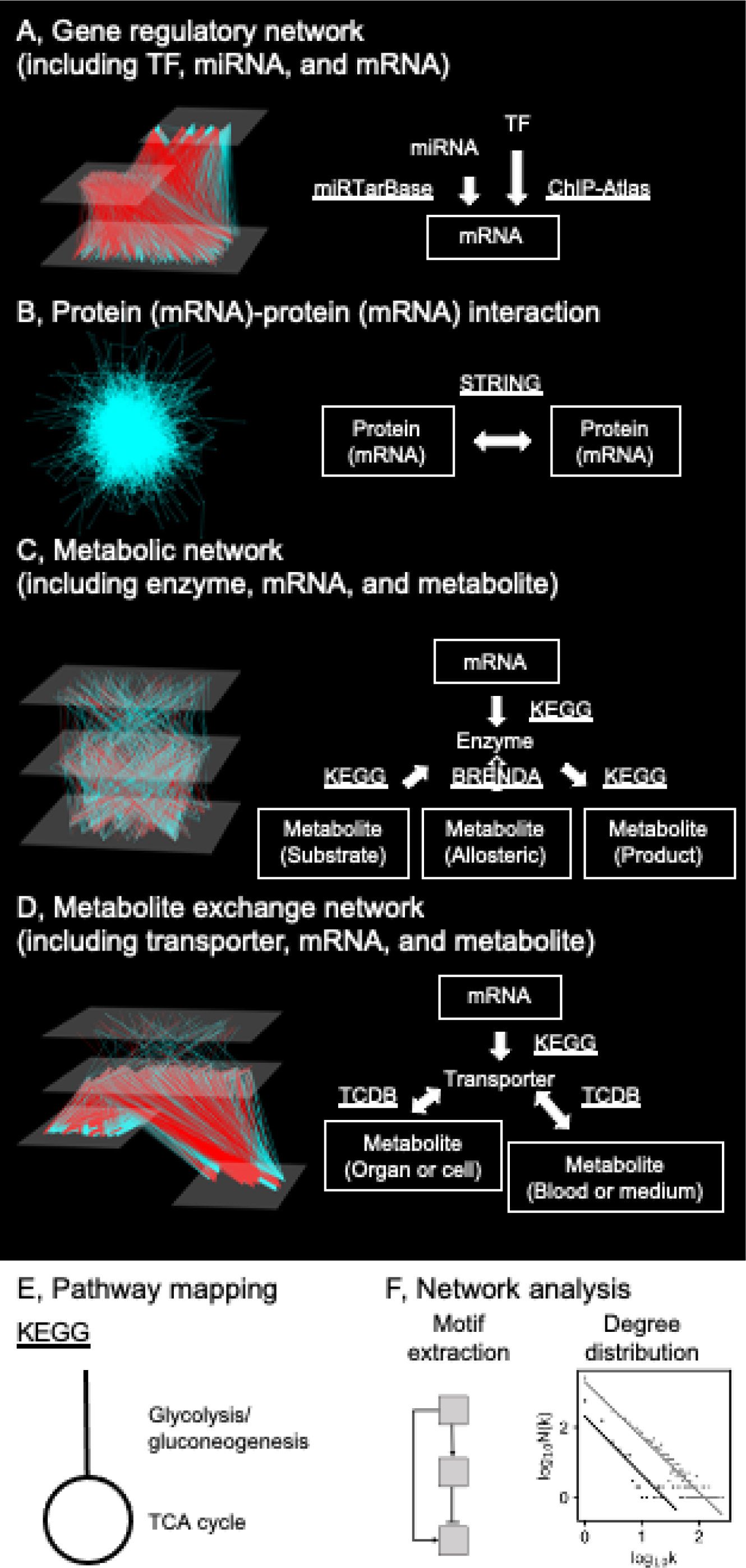
Outputs of iTraNet. (A) A network from transcription factor (TF) or microRNA (miRNA) to mRNA. TFs and miRNAs are estimated using the ChIP-Atlas and miRTarBase databases, respectively. (B) A network connecting mRNAs or proteins. The association is based on protein–protein interactions and is estimated by STRING. mRNAs can be used as proxies for proteins. (C) A network connecting mRNA, enzyme, and metabolite. The connections are estimated using KEGG and BRENDA. (D) A network connecting mRNA, transporter, and metabolite. The connections are estimated using KEGG and TCDB. (E) iTraNet visualizes trans-omics networks by arranging each metabolic pathway using the well-known KEGG layout. (F) iTraNet performs motif extraction from trans-omics networks and analyzes topological features of trans-omics networks, such as degree distributions. The parameter *N*(*k*) is the number of nodes in the network with *k* connections to other nodes.

iTraNet also estimates mRNAs or proteins associated with the uploaded mRNAs or proteins (Fig. 2B). The association is based on PPIs and is estimated using the STRING database (19). Only Mus musculus data in STRING was used in this web application. The network construction employs a cutoff score of 400, referencing the default value in STRING (19).

iTraNet also estimates metabolic reactions and enzymes associated with the uploaded mRNAs and metabolites (Fig. 2C). Metabolic reactions are assumed to be catalyzed by metabolic enzymes and affected by metabolites that function as the substrates, products, or allosteric regulators. Regulation of metabolic reactions also consists of regulations by changing the amount of enzyme through gene expression. This inference draws from analysis of data from Kyoto Encyclopedia of Genes and Genomes (KEGG) (20) and BRaunschweig ENzyme DAtabase (BRENDA) (21). These molecules are organized into a global metabolic pathway (mmu01100) in the KEGG database. Mus musculus data in KEGG database was used in iTraNet. If users only have proteome data and not transcriptome data, the proteome data can be converted from ENSMUSP to ENSMUSG using Ensembl BioMarts (22), etc., and uploaded as transcriptome data for this analysis.

iTraNet also estimates transporters associated with the uploaded mRNAs, metabolites in the target organ or cell, and metabolites in blood or medium data (Fig. 2D). Transporters are assumed to be affected by metabolites and regulations by changing the amount of transporter gene expression. This inference draws from analysis of data from the Transporter Classification Database (TCDB) (23). If users only have proteome data and not transcriptome data, the proteome data can also be converted from ENSMUSP to ENSMUSG and uploaded as transcriptome data for this analysis.

iTraNet also visualizes trans-omics networks by arranging each metabolic pathway, such as glycolysis/gluconeogenesis and the tricarboxylic acid (TCA) cycle using the well-known KEGG layout, thereby enhancing their interpretability (Fig. 2E). Of note, all network types described above are presented as interactive networks, enabling users to zoom in/out and drag nodes and edges as needed.

### Network analysis

To understand living systems, it is necessary not only to visualize and investigate how each molecule interacts but also to study the overall network properties of how all molecules interact, which goes beyond reductionist methods (1, 24). Moreover, network analysis identifying pivotal nodes, hubs, and modules within complex networks have guided hypothesis formation and developed targeted intervention strategies (7–10). To achieve such a comprehensive understanding of living systems, iTraNet also calculates properties of the trans-omics networks (Fig. 2F). In this web application, trans-omics networks are depicted as graphs where nodes represent various molecules, including mRNAs, miRNAs, proteins, or metabolites. Edges symbolize physical or functional interactions, following the typical representation (12, 25).

A key characteristic of biological networks is the majority-leaves minority-hubs (mLmH) topology (1, 25), and iTraNet investigates the topological features of trans-omics networks. Of note, mLmH-based networks are often referred to as scale-free networks (25); however, here, we employ the term mLmH to emphasize that this application neither assumes nor advocates the strict power–law distribution concept associated with scale-free networks, as it has generated some debate in the field (25). iTraNet calculates degree and degree centrality of each node, and then calculates the degree distributions of trans-omics networks. “Degree” refers to the number of connections or edges that a node has within the network. It indicates the level of connectivity of a node in relation to other nodes. “Degree centrality” is a measure that quantifies the importance of a node based on the number of its connections. It provides insights into the node’s influence within the network. “Degree distributions” denote the distribution of nodes based on their degrees, indicating the pattern of connectivity across the network. iTraNet also calculates a scaling parameter (*γ*), which is defined as follows:

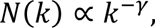

where *N*(*k*) denotes the number of nodes in the network with *k* connections to other nodes, and *γ* is estimated through least-squares fitting. Of note, commonly used methods for analyzing power–law distributions, such as least-squares fitting, can yield imprecise *γ* (26), and the value provided by iTraNet should be interpreted cautiously.

iTraNet also calculates several network metrics, including density, clustering coefficient, assortative, and the mean degree of trans-omics networks. “Density” refers to the ratio of the number of connected edges present in a network to the total number of possible edges. It provides an indication of how interconnected the nodes are within the network. “Clustering coefficient” quantifies the extent to which nodes in a network tend to form clusters or groups. It measures the probability that the neighbors of a node are also connected to each other. “Assortative” refers to a network characteristic that describes the tendency of nodes with similar degrees to be connected. Positive assortative indicates that nodes with high degrees tend to connect with other nodes with high degrees, and nodes with low degrees connect to nodes with low degrees. “Mean degree” represents the average number of edges connected to nodes in a network. It provides an overall view of the network’s connectivity.

These measures collectively indicate the structural properties and connectivity patterns within the trans-omics networks.

Biological networks can lead to nonlinearity in the input–output relationship when loops are formed, as exemplified by the incoherent feedforward loop (27). Such network motifs can be associated with intriguing biological phenomena. Hence, iTraNet also incorporates the capability to extract loop structures from trans-omics networks.

### Server implementation

The iTraNet web-based server is implemented in Python and uses the Streamlit library (https://www.streamlit.io) for the web application. iTraNet is hosted on the Streamlit cloud.

The stand-alone tool to run iTraNet on a local machine is also available at the GitHub repository (https://github.com/HikaruSugimoto/Transomics_iTraNet). This code can be freely modified in a local machine to add or modify visualization and analysis methods. Files are written using Pandas (28). Trans-omics networks were visualized and analyzed using Networkx (29), Matplotlib (30), and PyViz (an open-source visualization and analysis packages in Python https://pyviz.org/).

## RESULTS

In this section, we summarize an overview of the application’s capabilities. We also describe case studies that use this application to demonstrate the applicability of the web application.

### The iTraNet web server

iTraNet is an interactive web application developed using Streamlit (https://www.streamlit.io), and the running instance of the online app is accessible through the website (https://transomics.streamlit.app/). The application is also available for local use by using the code provided in the GitHub repository (https://github.com/HikaruSugimoto/Transomics_iTraNet). iTraNet is freely available to all users, and there is no login requirement. Users can directly upload their own data in a simple format that contains molecular IDs and type of responses (up or down).

iTraNet accepts transcriptome, proteome, and/or metabolome data as input. Additionally, users have the option to upload background genes when performing TF enrichment analysis. The output generated by iTraNet encompass four distinct types of network visualizations and analyses, as described in the Methods section: A, gene regulatory networks (including TF, miRNA, and mRNA) estimate TFs and miRNAs associated with the uploaded transcriptome data; B, protein (mRNA)–protein (mRNA) interactions estimate proteins or mRNAs associated with the uploaded proteome or transcriptome data; C, metabolic networks (including enzyme, mRNA, and metabolite) estimate metabolic reactions and enzymes associated with the uploaded transcriptome and metabolome data; and D, metabolite exchange networks (including transporter, mRNA, and metabolite) estimate transporters associated with the uploaded transcriptome and metabolome data.

### Case study 1: Comprehensive analysis of the metabolic status after glucose administration

To showcase the utility of iTraNet in visualizing and analyzing trans-omics networks, we employed publicly available transcriptome and metabolome data from the mouse liver, muscle, and blood following the oral glucose tolerance test (11, 12). In these datasets, 16- hour fasted wild-type (WT) and *ob/ob* mice were orally administered glucose. Livers, muscles, and blood at 0, 20, 60, 120, 240 min after glucose administration were collected (Fig. 3A). The mRNAs and metabolites that exhibited significant changes after glucose administration, as determined by statistical methods described in previous studies, were used as input for iTraNet. The number of differentially expressed molecules is depicted in Figure 3B.

**Figure 3.**
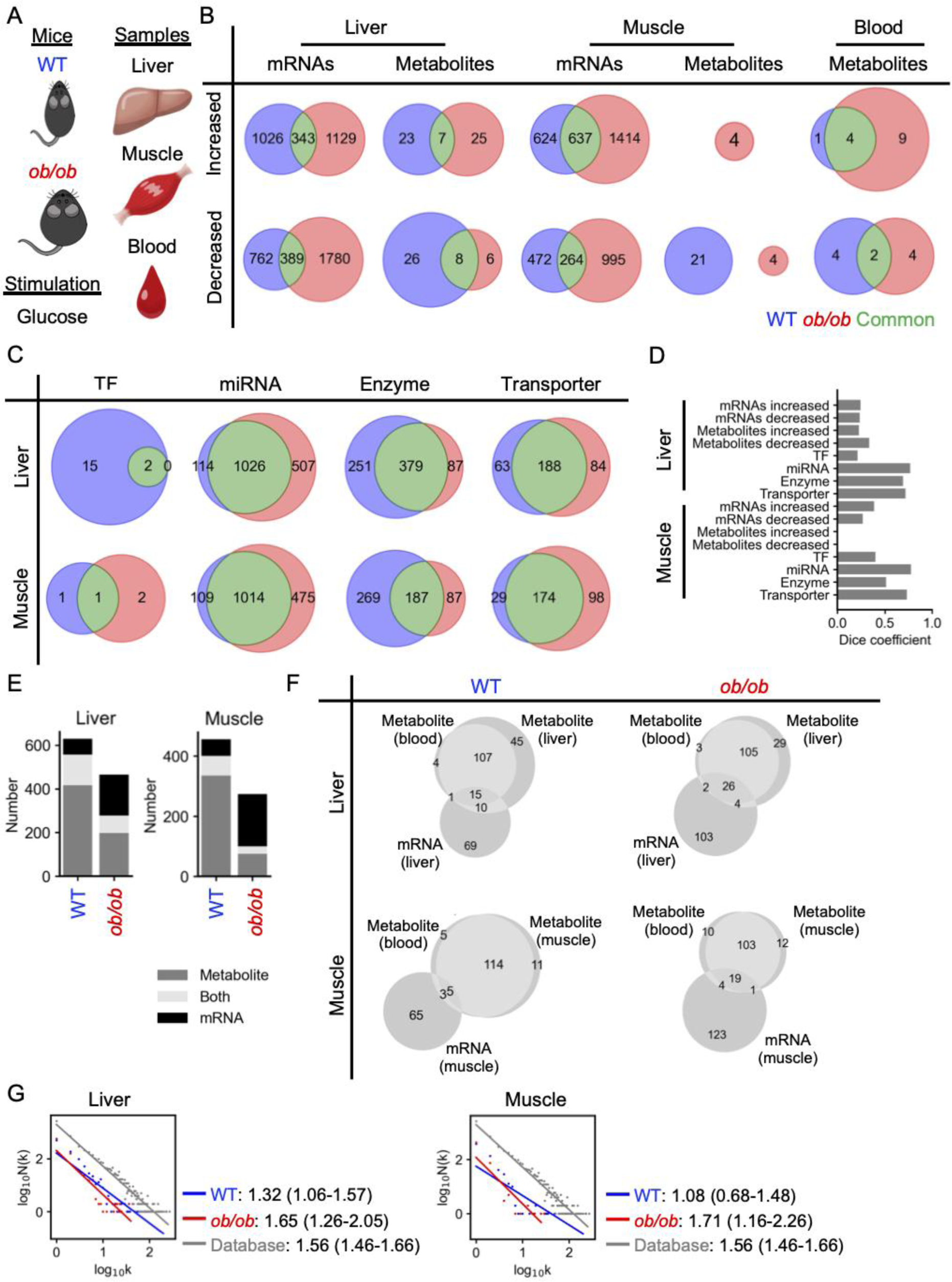
Comprehensive analysis of the metabolic status after glucose administration using iTraNet. (A) As a case study of iTraNet, we used previously reported transcriptome and metabolome data from WT and *ob/ob* mice liver, muscle, and blood after an oral glucose tolerance test. (B) The number of differentially expressed molecules in each condition. Molecules that exhibited changes specific to WT mice (blue), changes specific to *ob/ob* mice (red), and changes in common to both WT and *ob/ob* (green). (C) The number of differentially expressed transcription factors (TFs), miRNAs, enzymes, and transporters predicted by iTraNet in each condition. (D) The dice coefficient of each Venn diagram shown in Figures. 3B and 3C. A high value of the dice coefficient indicates the corresponding molecules are similar between the groups. (E) The number of predicted enzymes associated with the mRNAs, metabolites, or both in liver and muscle. (F) The number of predicted transporters associated with the mRNAs, metabolites, or both in liver and muscle. (G) Degree distributions with fitted regression lines for the trans-omics network. N(k) represents the number of nodes in the network with k connections to other nodes. Gray, the degree distribution for the network consisting of all regulations in the KEGG and BRENDA databases; blue, the degree distribution for the network consisting of only differentially expressed reactions in WT; red, the degree distribution for the network consisting of only differentially expressed reactions in *ob/ob*. The values are the scaling parameters of the degree distributions, and the values in the parenthesis are 95% confidence interval.

Upon input submission, iTraNet generated four types of biological networks and visualized the networks by arranging each metabolic pathway using the KEGG layout, as outlined in Figure 2. Since most visualization aspects have been previously discussed (11, 12), we mainly present the outcomes of molecular estimations and network analyses.

iTraNet also estimated TFs, miRNAs, enzymes, and transporters associated with the differentially expressed molecules (Fig. 3C). To investigate which molecule types exhibited significant differences between WT and *ob/ob*, we calculated the dice coefficient, which increases if the corresponding molecules are similar between the groups (Fig. 3D). While the dice coefficient for mRNAs in the liver and muscle were comparable, the coefficient for liver metabolites was higher than that for muscle metabolites, suggesting distinct regulation of muscle metabolites in *ob/ob*. In both the liver and muscle, the dice coefficients for enzymes and transporters were larger than those for mRNAs and metabolites. These results indicate that in *ob/ob*, the response of mRNAs and metabolites to glucose was relatively different from that in WT; however, these mRNAs and metabolites were not completely dissimilar; rather, specific molecules among multiple mRNAs and metabolites associated with particular enzymes or transporters exhibited differences between WT and *ob/ob*.

iTraNet further estimated the number of enzymes associated with the mRNAs, metabolites or both (Fig. 3E). In both the liver and muscle, more enzymes were influenced by mRNAs, while fewer enzymes were affected by metabolites in *ob/ob*, as previously described (11, 12). iTraNet also estimated the number of transporters associated with the mRNAs or metabolites or both (Fig. 3F). In both the liver and muscle, more transporters are affected by mRNA in *ob/ob*. Collectively, these results indicate that *ob/ob* exhibited an increased number of differentially expressed mRNAs affecting metabolic enzymes and transporters.

iTraNet also investigated the topological features of the trans-omics network. Among the analyses, degree distributions of the networks including enzyme, mRNA, and metabolite are shown in Figure 3G. The trans-omics network encompassing all regulations within the KEGG and BRENDA databases (Fig. 3G, gray line) can be considered a majority- leaves minority-hubs (mLmH) network with a power–law degree distribution. In both liver and muscle, the scaling parameters of the degree distribution of the WT network were smaller than those of the background network, indicating that WT tends to use hub nodes after glucose administration. By contrast, the scaling parameters of the degree distributions of the *ob/ob* networks were larger than those of the WT networks, indicating that hub nodes in WT tend to be unresponsive to glucose administration in *ob/ob*.

### Case study 2: Comprehensive analysis of the metabolic status under a cold environment

To further showcase the utility of iTraNet in visualizing and analyzing trans-omics networks, we employed another publicly available transcriptome and metabolome data (31). In this dataset, brown adipose tissue (BAT) and blood from mice at room temperature (23°C) or cold (4°C) for 5 hours were investigated (Fig. 4A). The mRNAs and metabolites were comprehensively examined by RNA sequencing (RNA-seq) and gas chromatography–mass spectrometry (MS), respectively. The mRNAs and metabolites that exhibited significant changes between the two groups, as determined by statistical methods described in the previous study, were used as input for iTraNet. The number of significantly increased and decreased molecules under the cold condition is depicted in Figure 4B.

**Figure 4.**
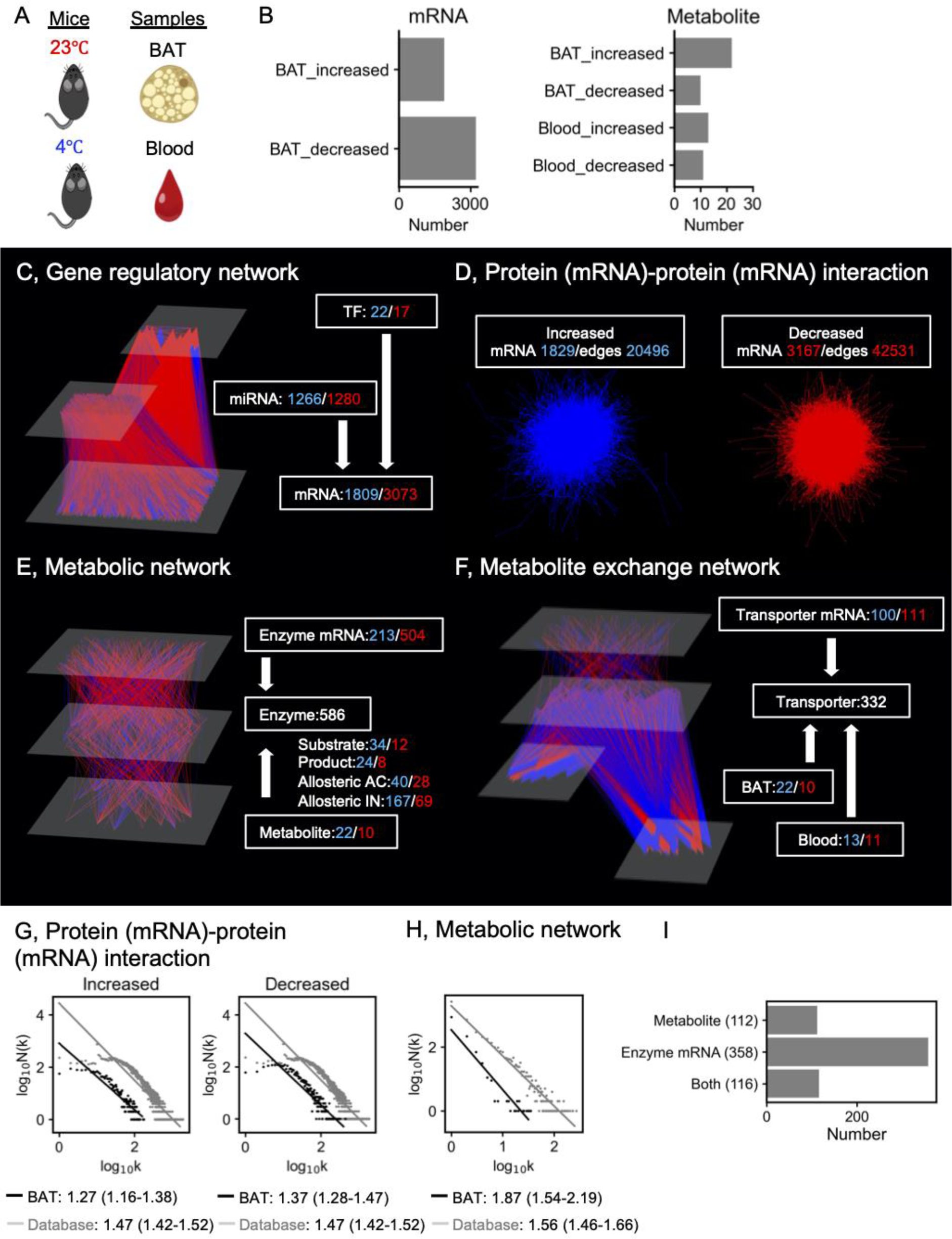
Comprehensive analysis of the metabolic status under the cold condition using iTraNet. (A) As a case study of iTraNet, we used previously reported transcriptome and metabolome data from brown adipose tissue (BAT) and blood from mice at room temperature (23°C) or cold (4°C). (B) The number of differentially expressed molecules in each condition. (C) A network connecting transcription factor (TF), microRNA (miRNA), and mRNA. TFs and miRNAs are estimated using iTraNet. Each number denotes the number of the associated molecules. Number with blue denotes the number of increased molecules under the cold condition, and that with red denotes the number of decreased molecules under the cold condition. (D) A network connecting mRNAs. The association is based on protein-protein interactions, and mRNAs are used as proxies for proteins. The blue network is created from mRNAs that increased in the cold environment, and the red one is created from mRNAs that decreased in the cold environment. (E) The trans-omics network for differentially regulated metabolic reactions. Nodes and edges indicate differentially expressed molecules and differential regulations, respectively. The Enzyme mRNA and Metabolite (Substrate, Product, Allosteric AC (activation), and Allosteric IN (inhibition)) layers correspond to differentially expressed mRNAs and differentially expressed metabolites, respectively. The Enzyme layer represents metabolic enzymes regulated by differentially expressed molecules. The differential regulations are classified into either activating (blue edges) or inhibiting (red edges). Increased Enzyme mRNA, increased Substrate, decreased Product, increased Allosteric AC, and decreased Allosteric IN are assumed to activate the reactions. (F) The trans-omics network for differentially regulated transporter. Nodes and edges indicate differentially expressed molecules and differential regulations, respectively. The Transporter mRNA and Metabolite (BAT and Blood) layers correspond to differentially expressed transcripts and differentially expressed metabolites, respectively. The Transporter layer represents transporters associated with differentially expressed molecules. The differential regulations are classified into either activating (edges with blue) or inhibiting (edges with red). Increased transporter mRNAs and increased metabolites are assumed to activate the reactions. (G) Degree distributions with fitted regression lines for the mRNA-mRNA network, which is based on protein-protein interactions shown in Figure 4D. N(k) represents the number of nodes in the network with k connections to other nodes. Gray, the degree distribution for the network consisting of all regulations in the STRING database; black, the degree distribution for the network consisting of only differentially expressed relation in BAT. The values are the scaling parameters of the degree distributions, and the values in the parenthesis are 95% confidence interval. (H) Degree distributions with fitted regression lines for the trans-omics network shown in Figure 4E. N(k) represents the number of nodes in the network with k connections to other nodes. Gray, the degree distribution for the network consisting of all regulations in the KEGG and BRENDA databases; black, the degree distribution for the network consisting of only differentially expressed reactions in BAT. The values are the scaling parameters of the degree distributions, and the values in the parenthesis are 95% confidence interval. (I) The number of predicted enzymes associated with the mRNAs or metabolites or both in BAT.

Upon input submission, iTraNet generated and visualized four types of biological networks (Fig. 4C–F). iTraNet estimated TFs and miRNAs associated with the uploaded mRNAs (Fig. 4C). Table 1A shows the TFs and miRNAs associated with many mRNAs. Among them, previously reported key molecules in BAT function, such as Pparg (TF), which regulates *Ucp1* (mRNA) (32), were included, suggesting some validity of the analysis.

**Table 1.**
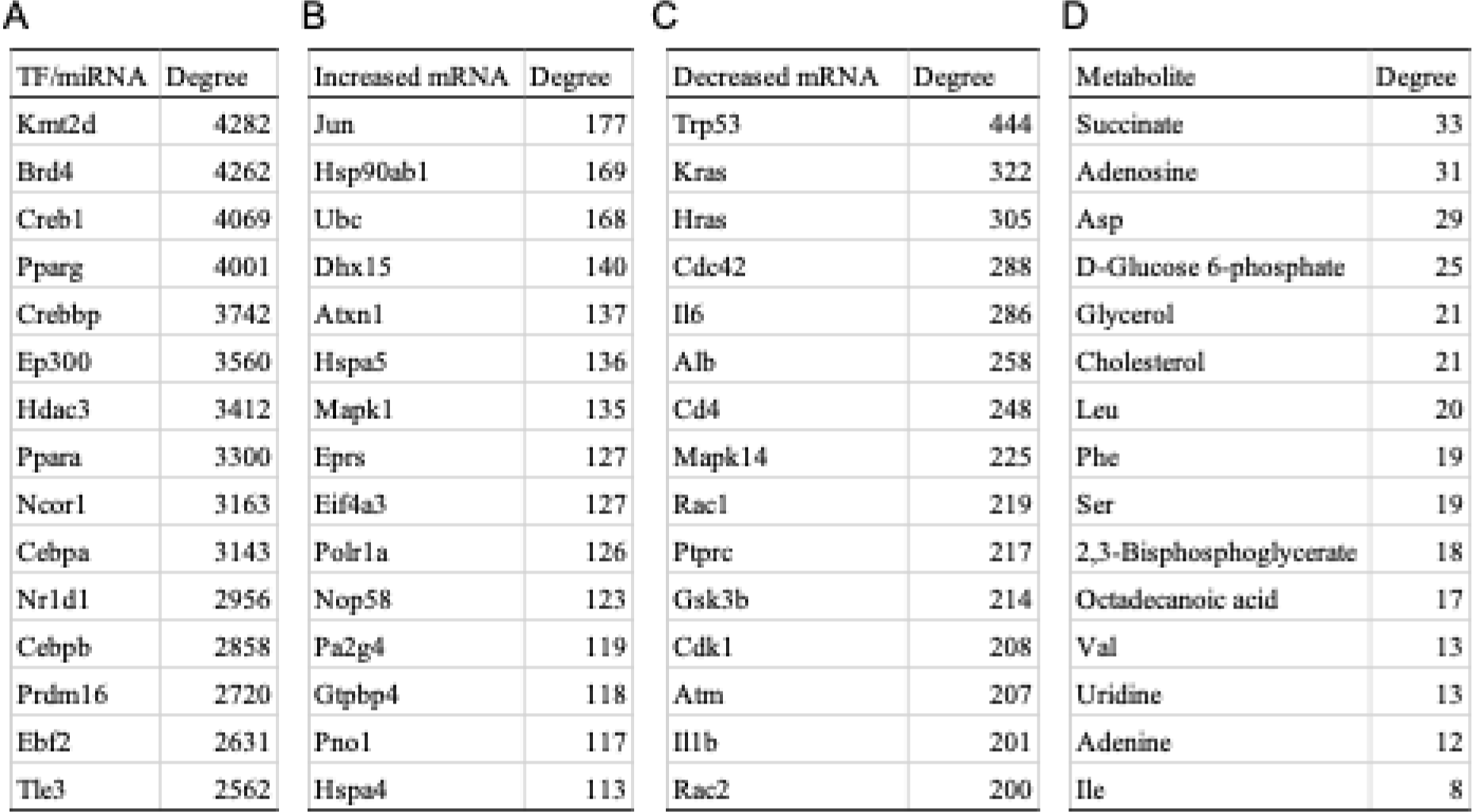
List of molecules with high degrees. List of 15 molecules with high degrees in the networks of. **Fig. 4C** **(A),** **Fig. 4D, increased (B),** **Fig. 4D, decreased (C), and** **Fig. 4E** **(D), respectively.**

iTraNet also estimated the protein (mRNA)–protein (mRNA) interactions (Fig. 4D) and investigated the topological features of the network (Fig. 4G). The protein (mRNA)– protein (mRNA) interaction network including all relationships within the STRING databases (Fig. 4G, gray line) can be considered a mLmH network with a power–law degree distribution. In both the increased and decreased networks, the scaling parameters of the degree distributions were smaller than those in the background network (Fig. 4G), indicating that mice tend to use hub mRNA nodes under the cold condition. By contrast, the scaling parameter of the enzyme, mRNA, and metabolite network was larger than that of the background network, indicating that mice did not tend to use the hub nodes of the enzymes, mRNAs, and metabolite network under the cold condition.

iTraNet also estimated the number of enzymes associated with the mRNAs, metabolites, or both (Fig. 4I). Although many metabolites including glucose and amino acid metabolism were significantly changed in their concentrations both after glucose loading (11, 12) and cold stimulation (31), many enzymes were found to be affected by metabolite changes after glucose loading (Fig. 3G) and by mRNA changes under cold stimulation in WT mice (Fig. 4I). These results collectively indicate that the tendency to use hub nodes of biological networks and mRNAs or metabolites in regulating metabolic reactions can vary by organ or condition. For a deeper understanding of metabolism, we can also use iTraNet to look up the names of the nodes in order of increasing degree in each network (Table 1).

## DISCUSSION

iTraNet stands as a user-friendly interactive web application designed for visualizing and analyzing trans-omics networks. This web application is capable of estimating TFs, miRNAs, enzymes, and transporters associated with transcriptome, proteome, and/or metabolome data uploaded by users. It also visualizes four distinct interactive biological networks: (A) gene regulatory networks (including TF, miRNA, and mRNA); (B) protein (mRNA)–protein (mRNA) interactions; (C) metabolic network (enzyme, mRNA, and metabolite); and (D) metabolite exchange networks (including transporter, mRNA, and metabolite). Additionally, the web application conducts analyses of network properties, exemplified by degree distributions as demonstrated in the case studies.

We used iTraNet to investigate the metabolic status after glucose loading (Fig. 3). In WT mice, hub molecules within the trans-omics network tended to respond to glucose administration, whereas this tendency disappeared in *ob/ob* mice. Metabolic networks have a mLmH topology, which is remarkably resilient to errors, with local errors rarely leading to a loss of global information transmission capacity within the network due to the presence of hub molecules (6). Therefore, disruption of homeostasis in biological networks with mLmH topology may inevitably involve changes in the configuration of the overall topology of the network, including changes in hub molecules, as observed in *ob/ob* mice. Of note, although the coverage of metabolites that can be measured by MS is high, it is less comprehensive than RNA-seq. Due to this coverage difference, the main focus in examining the effects of mRNAs and metabolites on enzymes will be on relative comparisons, such as the comparison between WT and *ob/ob* mice, as described in this study.

We also investigated the metabolic status under cold stimulation (Fig. 4) using iTraNet. Both glucose administration (11, 12) and cold stimulation (31) affect various metabolites, including glucose and amino acid metabolism, but the regulation of these molecules and the overall topology of the networks might be different between the two conditions (Figs. 3, 4). The tendency to use hub nodes in biological networks and the tendency to use either mRNAs or metabolites to regulate metabolic reactions varied among organs or conditions. Such a comprehensive understanding and overview of living systems requires comprehensive molecular measurements and trans-omics analysis, as facilitated by this web application. Network analyses have also identified pivotal nodes within complex networks and have guided hypothesis generation (7–10). For this purpose, iTraNet can also output the name of each hub node, as shown in Table 1, which includes previously reported key molecules in BAT function. Collectively, we expect that iTraNet can facilitate researchers in understanding biological dynamics both from the perspective of the overall network properties and from the perspective of individual molecular interactions.

Various web applications have been developed for visualizing and analyzing multiomics data such as OmicsNet (2, 33), 3Omics (34), PaintOmics (35), MergeOmics (36), MiBiOmics (37), OmicsAnalyst (38), and Arena3D (39, 40). The detailed characteristics of these web-based tools were well summarized and compared previously (2, 38). Compared to conventional web applications such as OmicsNet, one of iTraNet’s distinct strengths lies in its capacity to arrange multiomics molecules within each metabolic pathway, such as glycolysis/gluconeogenesis and the TCA cycle, using the well-known KEGG layout. This feature is anticipated to facilitate a more intuitive understanding of intricate biological networks by researchers. Moreover, iTraNet can estimate more various regulations, such as allosteric regulation and regulation by transporters. We expect that this web application can deepen the understanding of biological phenomena by visualizing individual molecules on the well-known metabolic map and simultaneously analyzing the overall features of the trans-omics networks that include more various regulations.

The current web server has several limitations. Due to storage constraints, iTraNet is currently designed exclusively for Mus musculus data. However, it is possible to apply the code in this web application to other species while maintaining the same framework, and the code in the GitHub repository (https://github.com/HikaruSugimoto/Transomics_iTraNet) can be downloaded and modified to analyze other species data. Of note, the KEGG database requires a license for non-academic users (https://www.pathway.jp/en/licensing.html).

Moreover, the web application is provided with limited memory, which might lead to slower access and computations. For faster and more stable calculations, users can download the code from the GitHub repository (https://github.com/HikaruSugimoto/Transomics_iTraNet) and run iTraNet on a local machine. iTraNet takes lists of mRNAs and metabolites as input, and the output of RNA-seq and MS cannot be directly used as the input; however, transcriptome and metabolome data can be analyzed using other well designed web servers, such as iDEP (41) and MetaboAnalyst (42). Furthermore, this web application still estimates a limited kind of molecular interactions. Future development of a web application that includes other molecular interactions, such as DNA methylation, long non-coding RNA, and phosphorylation is warranted.

In conclusion, here we introduced iTraNet, a web application that provides users with visualizations and analyses of trans-omics networks. Given its ability to easily visualize the intricate interactions of the various biological molecules and perform trans-omics network analyses, we expect that it will facilitate researchers in understanding complex biological dynamics.

## Acknowledgments

We thank Katsuyuki Yugi and Kozo Nishida for their assistance with the analysis and critical reading of this manuscript.

## Contribution statement

H.S. conceived the project. H.S., K.M., D.L., and Y.B. analyzed the data. H.S., K.M., D.L., Y.B., M.M., and S.K. wrote the manuscript. S.K. supervised the study.

## Funding and assistance

This study was supported by the Japan Society for the Promotion of Science (JSPS) KAKENHI (JP21H04759), CREST, the Japan Science and Technology Agency (JST) (JPMJCR2123), and The Uehara Memorial Foundation.

## Data availability

iTraNet is freely available for all users at (https://transomics.streamlit.app/), and the source code is available at the GitHub repository (https://github.com/HikaruSugimoto/Transomics_iTraNet). The stand-alone tool to run iTraNet on a local machine is also available at the GitHub repository (https://github.com/HikaruSugimoto/Transomics_iTraNet).

